# Inverted Repeats in Viral Genomes

**DOI:** 10.1101/2025.11.10.687097

**Authors:** Madhura Sen, George M. Rivera, Jingxiang Gao, Matthew Shtrahman, Madhavi Ganapathiraju

## Abstract

An inverted repeat (IR) in DNA is a sequence of nucleotides that is followed by its complementary bases but in reverse order, occurring on the same strand (e.g., TCACCGCGGTGA). If the two complementary sequences occur one after the other without other bases between them, they are referred to as DNA palindromes. IRs could form hairpin and cruciform secondary structures, which endanger genomic stability. They are found to be prevalent in viral DNA at origins of replication, and they play a crucial role in various biological processes including gene silencing, duplication, and genomic evolution. IRs have been less explored, which stems from the scarcity of sequence analysis tools allowing accurate detection on large viral genome data. Here, using the Biological Language Modeling Toolkit (BLMT), we analyzed 14 thousand viral genomes for occurrences of IRs, resulting in the identification of over 19 million IRs longer than 20 bases, including 134 IRs that are 2000 bases long, and around 1,300 IRs per virus.

## Introduction

An inverted repeat is a sequence on a DNA strand that is followed by its complement appearing in reverse order on the same strand (1). The reverse complement may follow immediately after, in which case the pair are referred as a palindrome (e.g., CGAGCTCG) (2), or be separated by other bases, in which case it is called an inverted repeat (e.g., CGAGtctaCTCG) (3). Palindromes and inverted repeats where there are a few mismatches in base pairing are known as nearly perfect palindromes and inverted repeats (4). In this study, we discuss the role of inverted repeats (IR) and their mechanisms related to viral infections, explore existing computational tools for locating such patterns in a gene sequence while introducing a novel tool in Biological Language Modelling Toolkit (BLMT) (48), and subsequently present analysis on a large-scale of 14 thousand viral genomes determining IR density, lengths, percentile positions, and large terminal repeats across different viral hosts.

IRs have a propensity to form hairpin and cruciform structures (7,8), which interfere with DNA replication, DNA damage response, and other genetic mechanisms, induce chromosomal breakage and DNA rearrangements, leading to a variety of diseases, mainly viral infections (5,6). Human SARS-CoV-2 genome was shown to have an abundance of IRs compared to bat CoVs and and deemed to be associated with mutations and recombination events among CoV genomes during the infection phase (9). Likewise, 2022 monkeypox virus was also noted to differ from the 2018/2019 strain by around 50 variants with a mutation rate of 6 to 12 times higher than anticipated (10) and linked to IR sequences (11).

IRs are noted to associate with human genetic disorders such as X-linked congenital hypertrichosis syndrome and palindrome-mediated t(3;8) hereditary renal cell carcinoma (12–20). Palindromes were found to be involved in viral packaging, replication, and defense mechanisms (21–23), and are associated with replication initiation sites in viruses such as herpes simplex, varicella-zoster, Epstein-Barr virus (EBV), cytomegalovirus, and baculovirus, and with the recombination process in coronavirus (24–26). Both perfect or nearly perfect palindromes play significant roles in replication initiation and recombination events in viral genomes (27–30) and are associated with diseases including Grave’s disease, multiple sclerosis, and myasthenia gravis. IR-mediated hairpins/cruciforms are hypothesized to modulate replication or recombination; however, for foamy viruses specifically there is no established causal link between IR burden and human neurodegenerative disease, and the evidence is largely observational (31).

Understanding inverted repeats and their role in genetics requires the development of efficient methods to detect them. However, a major caveat arises wherever imperfect inverted repeats are involved, owing to the few mismatches in base pairs at the center or the gap sequence in between the complementary sequences. In this study, we employed the tools available in Biological Language Modelling Toolkit (BLMT) to find IRs using a novel approach. By concatenating each genome with its reverse complement and then using BLMT’s augmented suffix arrays to exhaustively find repeat pairs that occur once in the forward half and once in the reverse-complement half (thereby tagging the two arms of an IR), we were able to capture all the IRs found by the other tools (DetectIR, EMBOSS Palindrome, Palindrome Analyser, and IUPACpal), and also found significantly more IRs uniquely with this method.

## Data and Methods

### Data

#### Data for pilot study

Genome sequences of five viruses were collected from the NCBI (https://ftp.ncbi.nlm.nih.gov/genomes/Viruses/): ADE virus (ADE), adeno-associated virus (AAV), zika virus, southern bean mosaic virus (SBM), and soybean chlorotic blotch virus (SBC).

#### Data for comprehensive analysis of viral genomes

13 thousand fully sequence viral genomes were collected from the NCBI database, retrieved on February 1st, 2023 (https://ftp.ncbi.nlm.nih.gov/refseq/release/viral/viral.1.1.genomic.fna.gz).

### Methods

Suffix array is a data structure that stores suffixes of a sequence starting at every position in a lexicographic order. A suffix array can be augmented with longest common prefix array and rank array which are numerical arrays that allow navigating and pattern mining over suffix array computationally efficient. Biological language modeling toolkit (BLMT) was developed to process genome or proteome sequences into augmented suffix arrays, and to carry out various pattern search operations on it, including finding of k-mer/n-gram counts, location and length of long repeated sequences, and palindromes (i.e., where the reverse complements are not separated by gap sequences). Here, to find inverted repeats, the algorithm uses the observation about the nature of the two halves of IRs. Say, the two halves of the IR are represented as S and R corresponding to the first half sequence and its reverse complement; e.g., if the IR is CGAGtctaCTCG, S would correspond to CGAG and R would correspond to CTCG. If the complement strand of the DNA is considered from its 5’ to 3’, R’s complement would be CGAG, which is the same as S in the original sequence. Thus, if we concatenate the genome and its reverse complement, the long concatenated string will contain these four sequences in order—S, R, S′, and R′—with S′ and R′ appearing in the reverse-complement half. Thus, BLMT, which has a tool to find repeating sequences can be used to locate S and S’ (or R and R’) indicating that the positions corresponding to them are inverse repeats.

If the DNA sequence is AAACAGACACGTCTGAAA, the complement strand of the sequence will be TTTCAGACGTGTCTGTTT. The sequence with its reverse complement concatenated to it is AAACAGACACGTCTGAAATTTCAGACGTGTCTGTTT, which will be processed by BLMT. The repetitions found are in bold: AAA**CAGA**CACG**TCTG**AAATTT**CAGA**CGTG**TCTG**TTT. Here, the repetitions are between the main and complementary sequences, and the presence of an inverted repeat has been confirmed.

We first carried out a pilot study to evaluate the performance of the Biological Language Modelling Toolkit (BLMT) by comparing it to the four existing tools and the inverted repeats identified by them. Taking advantage of the efficiency of BLMT inferred from preliminary investigation, we employed BLMT to identify inverted repeats in all existing viral genomes on NCBI.

### Pilot study to evaluate computing IRs with BLMT

Five genomes collected for pilot study were run through the BLMT as described above. For each virus, we took the reverse complement of the entire sequence and concatenated it with the original gene sequence. We call them forback sequences (forward-back) and those were the input sequences for BLMT large repeat finding program (to indirectly find inverted repeats).

We also computed the IRs with the computational tools selected for comparison, namely: DetectIR (49), EMBOSS Palindrome (50), Palindrome Analyser (51), IUPACpal (52).

Length of each input sequence (in bp; the length of the genome plus its reverse compliment concatamer): ADE=29254, AAV=9534, Zika=21588, SBC=5416, SBM=8264

To give a consistent scope for all, we evaluated them with the same parameters. Parameters set for all programs and all sequences:

Maximum length of half-sequence = 7

Minimum length of half-sequence = 100

Maximum length of gap sequence = 100

Number of mismatches allowed = 0

### Study of All Viral Genomes Using BLMT

For each of the thirteen thousand fully sequence viral genomes from the NCBI database, we concatenated the genome sequence with its complementary sequence read in reverse order, which forms the input to the analysis. Suffix arrays, longest common prefix arrays and rank arrays were constructed for each input using BLMT. With find large repeats program from BLMT, we identified all repeats that are >= 9 bases. If a pair of repeats occurred at positions x_1_, y_1_, we verified whether y_1_ occurred after the midpoint of the new sequence, because it would then correspond to the second half of the IR in the main genome. By assessing whether it is a large repeat within the genome or whether it is a repeat between main and complementary strands, it can be determined whether it is an IR and if so, where it appears in the genome.

After performing the analysis for all viruses, we perform the same analysis for viruses on different hosts. Using the NCBI Virus Interface, we selected the six most important host categories: bacteria, fungi, humans, invertebrates, land plants, and vertebrates. For each host selected, we downloaded a complete RefSeq release of viral genomes for that host, and we performed the analysis following the same procedures.

## Results and Discussion

Figure 1 presents the total number of inverted repeat sequences found by each tool for each viral genome. It is clear that BLMT has discovered the highest number of sequences for all five viral genomes. The second highest number of sequences were detected by IUPACpal in ADE and AAV and by DetectIR in Zika, SBC and SBM.

**Figure 1:**
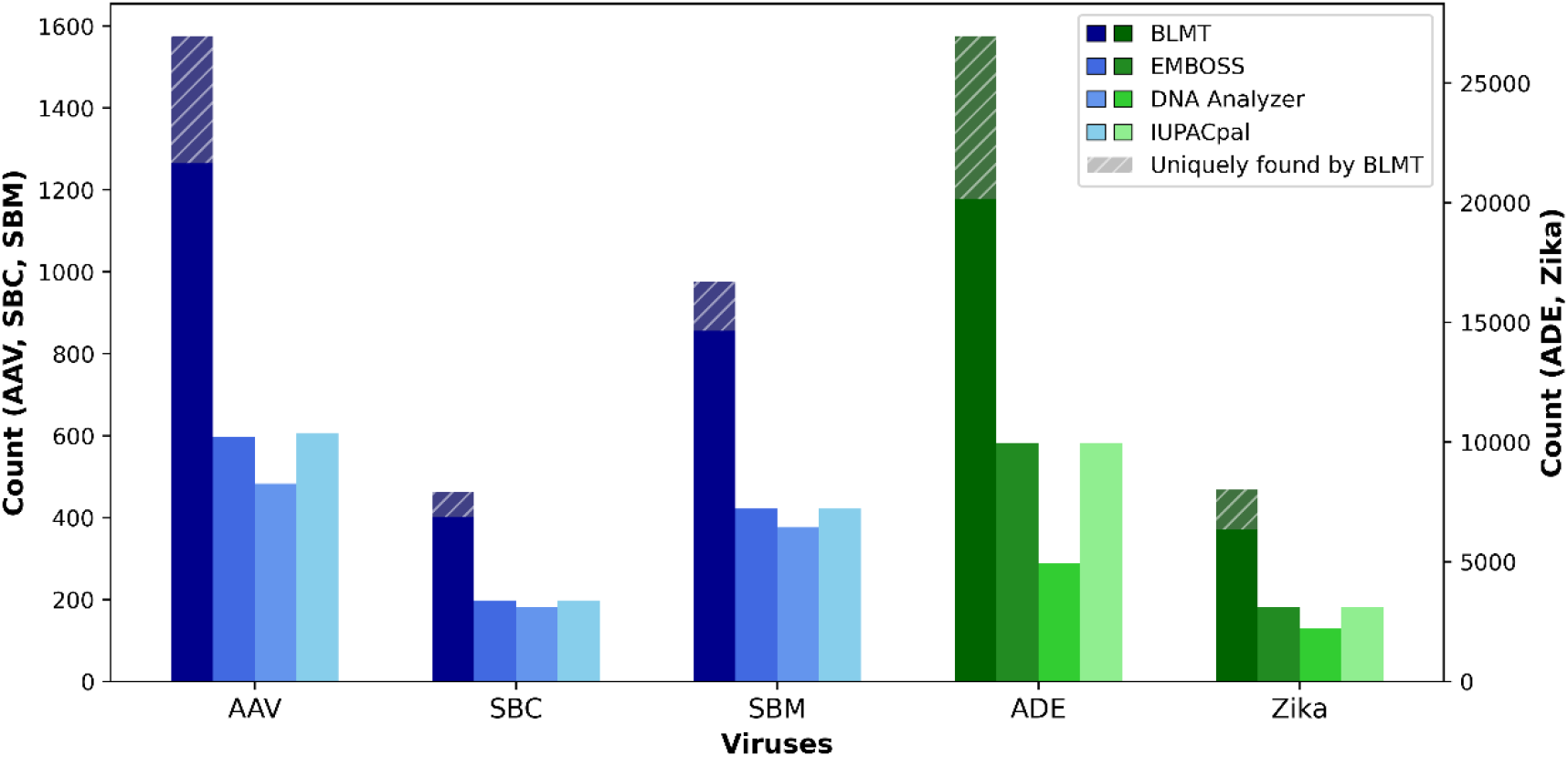
Comparison of IR identification by various tools.

We carried out the analysis of the complete RefSeq release of viral genomes from the NCBI database, which consisted of 14,797 genomes. We excluded files that contain unresolved nucleotide bases (represented as ‘N’ for any nucleotide), where the palindromic nature or complementarity of the two halves is ambiguous. After preserving files whose sequences are represented by A, T, C, and G, we have 13,023 files remaining for analysis.

### Lengths of inverted repeats

We first analyzed the mean and standard deviation of inverted repeats. Overall, the mean of length of inverted repeats is 10.6, with standard deviation 16.9. The distribution of mean lengths are as shown in Figure 2. The mean length of inverted repeats for the majority of viruses lies between 10 to 12. Table 1 shows the top 10 viruses with highest mean inverted repeat length.

**Table 1:** Viruses with highest average length of inverted repeats.

| ID | Virus Name | Average IR length |
| --- | --- | --- |
| 1 | *Fusarium graminearum mycovirus* | 108.5 |
| 2 | *Dendrolimus punctatus densovirus* | 42.5 |
| 3 | *Goose parvovirus* | 38.8 |
| 4 | *Planaria asexual strain-specific virus-like element type 1* | 34.4 |
| 5 | *Diatraea saccharalis densovirus* | 28.9 |
| 6 | *Muscovy duck parvovirus* | 27.7 |
| 7 | *Arracacha virus B strain* | 26.9 |
| 8 | *Aedes albopictus densovirus* | 24.8 |
| 9 | *Feldmannia irregularis virus a strain* | 24.4 |
| 10 | *Galleria mellonella densovirus* | 24.3 |

**Figure 2.**
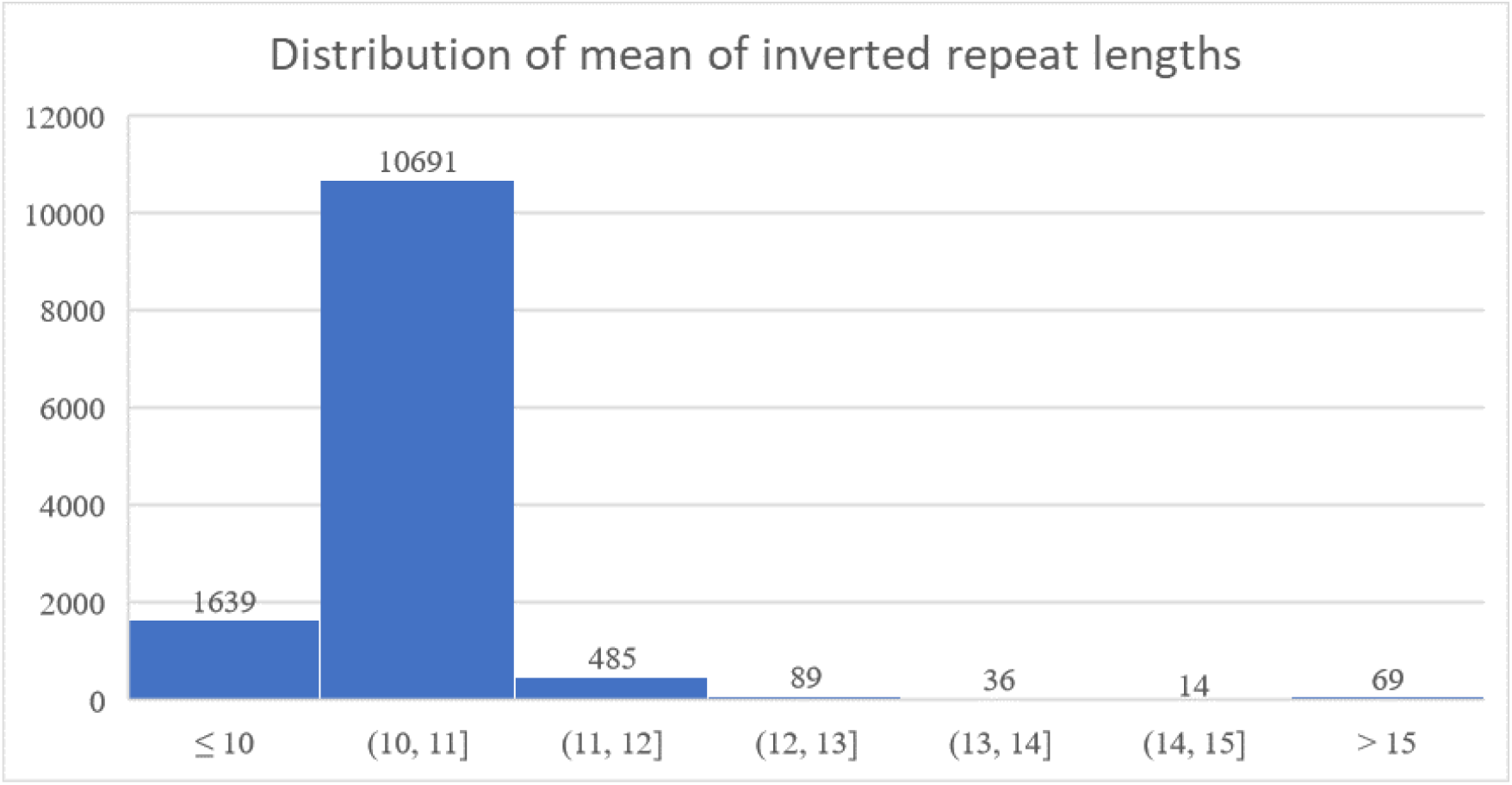
Distribution of mean of length of inverted repeats.

Among these viruses, there are many species from the *Densovirinae* subfamily, such as *Dendrolimus punctatus densovirus* and *Diatraea saccharalis densovirus*. Moreover, species of *Parvoviridae* such as *Goose parvovirus* and *Muscovy duck parvovirus* also have a high average length of inverted repeats. Each parvorirus contains linear, single-stranded DNA genomes with approximately 5 kb in size and encodes non-structural proteins, which are crucial for viral gene expression and replication (32). In densovirus, the replication stems from inverted terminal repeats present in the viral genome and involves a mechanism known as rolling circle replication similar to parvovirus (33,34).

Among these viruses, there are many species from the Herpesviridae subfamily, such as Gallid herpesvirus and Human herpesvirus. The large standard deviation indicates a wide dispersion of the lengths of inverted repeats, implying the existence of very large inverted repeats in these viruses. Some species of Poxviridae are also present, such as Racoon poxvirus. A study on herpes simplex virus 1 (HSV-1) concluded that replication within the genome occurs between the inverted repeats, producing concatemeric molecules reaching lengths equivalent to 10 times the viral genome (35). The current MPXV genomes were also identified to be replete with IRs ranging from 6754 to 8933, with a frequency of 34.25–45.11 per kbp (11).

### Normalized frequency of inverted repeats

The number of viruses with normalized frequency of inverted repeats for every 10,000 base pairs is shown in Figure 3.

**Figure 3.**
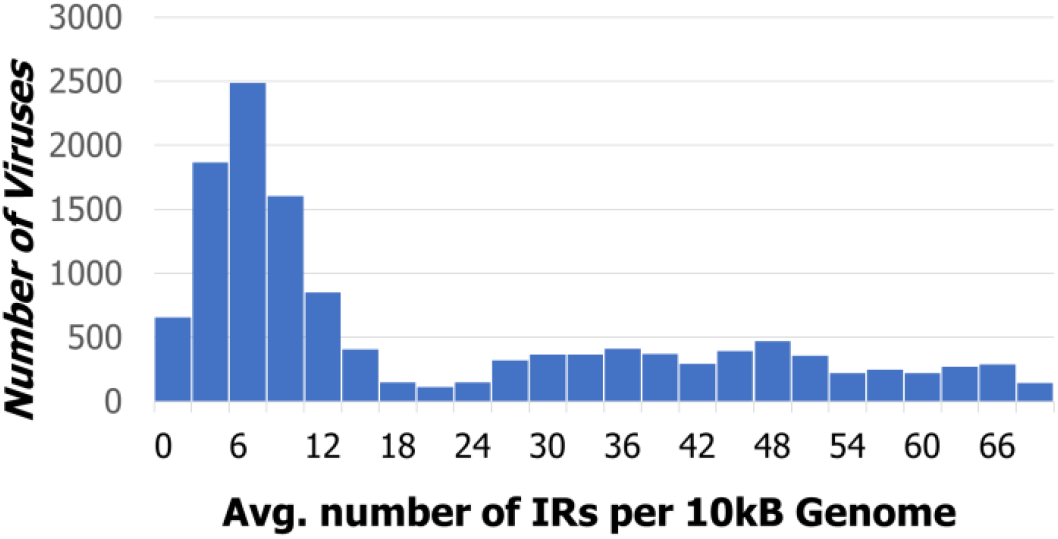
Distribution of viruses by number of inverted repeats per 10kb of genome length.

Overall, the mean normalized frequency is 20.0. Over half of the viruses having fewer than 15 inverted repeats per 10,000 base pairs. Tables 5 and 6 show the 10 viruses that have the lowest and highest normalized IR frequencies.

**Table 5:** Viruses with low normalized frequency of inverted repeats.

| ID | Virus name | Normalized Frequency (per 10k bps) |
| --- | --- | --- |
| 1 | Peanut stunt virus | 0.34 |
| 2 | Tasmanian aquabirnavirus | 0.34 |
| 3 | Bdellovibrio phage | 0.44 |
| 4 | Wuhan Mosquito Virus | 0.45 |
| 5 | Microviridae Fen7918 | 0.48 |
| 6 | Sclerotinia sclerotiorum mycoreovirus | 0.48 |
| 7 | Mushroom bacilliform virus | 0.50 |
| 8 | Syngnathus scovelli chapparvovirus | 0.50 |
| 9 | Southern rice black-streaked dwarf virus | 0.52 |
| 10 | Flammulina velutipes browning virus | 0.52 |

**Table 6:** Viruses with high normalized frequency of inverted repeats.

| ID | Virus name | Normalized Frequency (per 10k bps) |
| --- | --- | --- |
| 1 | Tenacibaculum phage | 77.81 |
| 2 | Bacillus phage | 76.90 |
| 3 | Lymphocystis disease virus | 75.15 |
| 4 | Fusobacterium phage | 73.78 |
| 5 | Yoka poxvirus | 73.78 |
| 6 | Acinetobacter phage | 73.39 |
| 7 | Campylobacter phage | 73.24 |
| 8 | Invertebrate iridescent virus | 72.86 |
| 9 | Turkeypox virus | 72.29 |
| 10 | Clostridium phage | 72.25 |

From Table 5, we can identify many fungi-related viruses with low IR frequency: Peanut stunt virus, Sclerotinia sclerotiorum mycoreovirus, Mushroom bacilliform virus, and Flammulina velutipes browning virus. Most fungal viruses with double-stranded RNA (dsRNA) have segmented genomes, which means their genetic material is divided into multiple segments, with each segment being enclosed in a different capsid (36). This structure inhibits the formation of long stretches of IRs since each segment contains specific genetic information (37). Also, fungal viruses employ replication strategies such as RNA-dependent RNA polymerase (RdRp)-mediated replication and reverse transcription that may not heavily depend on inverted repeats for effective viral replication, resulting in a lower prevalence of such sequences in their genomes (38,39).

Among viruses with high normalized IR frequencies, bacteriophage and poxvirus species commonly appear. Short inverted terminal repeats were discovered in small Bacillus bacteriophage genomes. The study investigated four phages (phi 15, Nf, M2Y, and GA-1) and determined the following terminal repeats, respectively: 5’A-A-A-G-T-A, 5’ A-A-A-G-T-A-A-G for Nf and M2Y, and 5’ A-A-A-T-A-G-A (40). Similarly, poxviruses contain ITRs with lengths ranging from 1 kb to >17 kb (41).

### IR positions within the Genome

We analyzed the positions at which inverted repeats start and end in all viral genomes, results of which are shown in Figure 4. Overall, inverted repeats seem to be more common in the 3’ half of viral genomes compared to 5’ half; there are fewer inverted repeats at the start or end of the viral genome compared to the middle.

**Figure 4.**
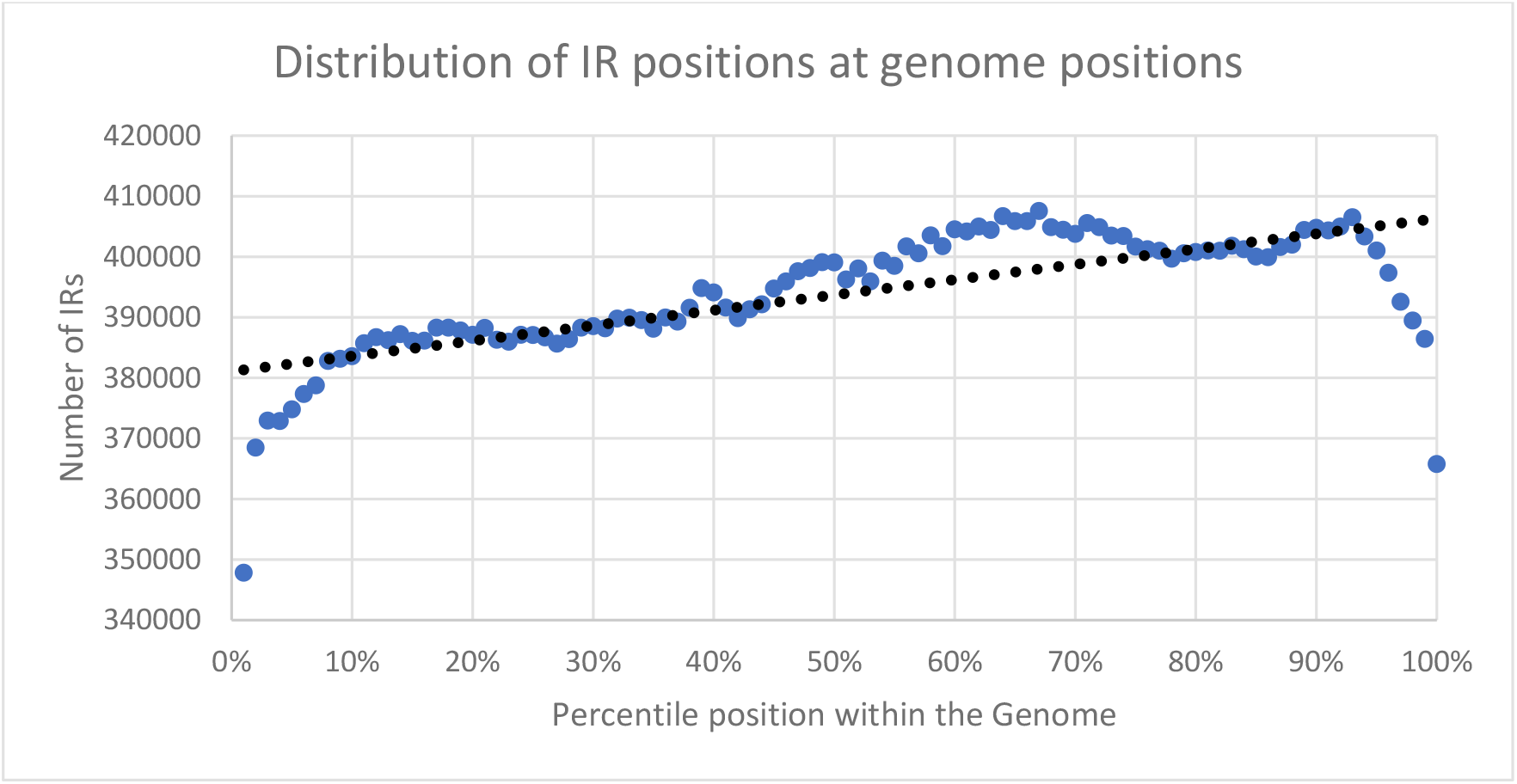
Distribution of percentile position of inverted repeats within viral genomes.

### Correlation between inverted repeats and viral hosts

We investigated the correlation between inverted repeats in viruses and the types of hosts they infect. After data collection and filtering, we have 4,567 virus genomes with bacteria hosts, 494 virus genomes with fungi hosts, 826 virus genomes with human hosts, 2,100 virus genomes with invertebrate hosts, 1,200 virus genomes with land plant hosts, and 2,676 virus genomes with vertebrate hosts.

We computed the mean lengths and normalized frequencies of inverted repeats for viral genomes in each of the categories, results of which are shown in Figure 5. The average inverted repeat length is similar for all virus categories, with a value between 10 and 11. However, the standard deviation for inverted repeat lengths is especially high for viruses with human, invertebrate, and vertebrate hosts, while it is very low for viruses with bacteria, fungi, and land plant hosts. We have listed the 10 human viruses with the highest average lengths in Table 7, and highest frequencies in Table 8.

**Table 7:** Human viruses with high average length of inverted repeats.

| ID | Virus name | Mean IR length |
| --- | --- | --- |
| 1 | Adeno-associated virus | 20.9 |
| 2 | Human parvovirus | 19.6 |
| 3 | Cyclovirus | 16.0 |
| 4 | Papiine herpesvirus | 14.2 |
| 5 | Macacine herpesvirus | 13.9 |
| 6 | Human herpesvirus | 13.6 |
| 7 | Puumala virus | 13.5 |
| 8 | Gyrovirus | 13.33333 |
| 9 | Tula virus | 13.33333 |
| 10 | Pulau reovirus | 13 |

**Table 8:** Human viruses with high normalized frequency of inverted repeats.

| ID | Virus name | IR normalized frequency (per 10k) |
| --- | --- | --- |
| 1 | Yaba-like disease virus | 70.4 |
| 2 | Variola virus | 70.0 |
| 3 | Akhmeta virus | 68.4 |
| 4 | Horsepox virus | 68.2 |
| 5 | Monkeypox virus | 67.8 |
| 6 | Molluscum contagiosum virus | 67.8 |
| 7 | Camelpox virus | 67.6 |
| 8 | Cowpox virus | 67.5 |
| 9 | Bovine papular stomatitis virus | 67.3 |
| 10 | Ectromelia virus | 67.1 |

**Table 10:** Table of viruses with long inverted terminal repeats.

| ID | Virus name | ITR length |
| --- | --- | --- |
| 1 | Raccoonpox virus | 18,952 |
| 2 | Cotia virus | 13,717 |
| 3 | Choristoneura rosaceana entomopoxvirus | 13,406 |
| 4 | Amsacta moorei entomopoxvirus | 9,457 |
| 5 | Horsepox virus | 7,526 |
| 6 | Mythimna separata entomopoxvirus | 7,346 |
| 7 | Canarypox virus | 6,490 |
| 8 | Monkeypox virus | 6,421 |
| 9 | Penguinpox virus | 5,765 |
| 10 | Taterapox virus | 4,778 |

**Figure 5:**
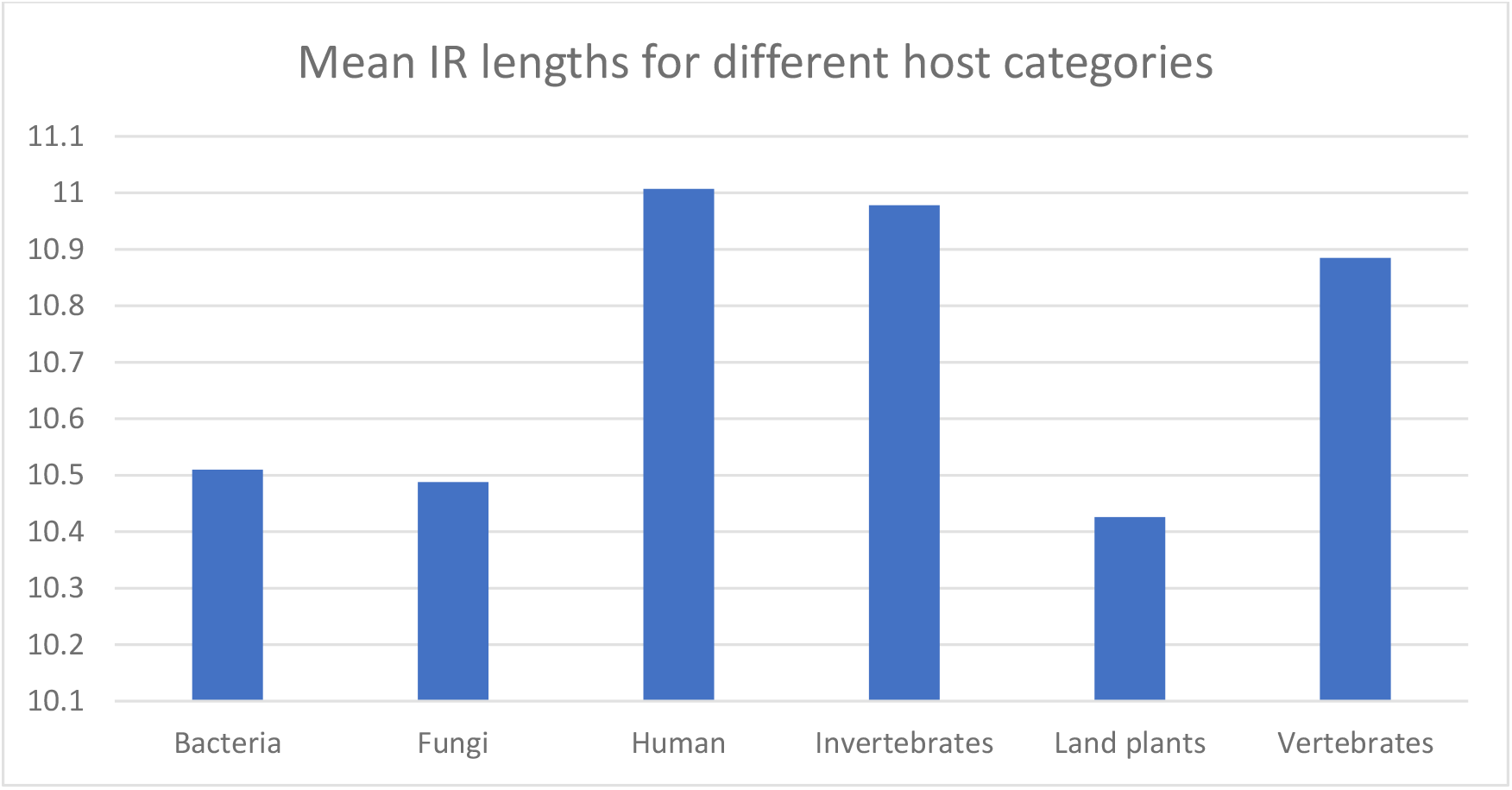
Mean and standard deviation of inverted repeat lengths for different virus categories.

Besides commonly occurring species of herpesvirus, direct and inverted repeats in cycloviruses were determined within their putative intergenic region in the genome (42). On the contrary, the IRs found in Puumala hantavirus were found in the 3’-noncoding region of the S segment. These IRs were found to play a role in recombination events that resulted in the deletion of the sequences responsible for forming hairpin structures (43).

The complete genome of Smallpox variola was also analyzed in other studies and found to contain 725 bp of ITRs with three 69-bp direct repeats. These terminal regions have numerous unique proteins with the potential to enhance the ability of the variola virus to spread and increase its virulence, specifically in humans (44). Akhmeta virus (AKMV), however, is a novel species of orthopoxvirus (OPXV). A genomic analysis demonstrated that AKMV also shared similarity with OPXV, which has approximately 6 kb sequence in the terminal region (45). Similar to the studies presented, tandem repeats were observed in Molluscum contagiosum virus, with the proponents associating the presence of such patterns with viral replication (46). Meanwhile, two identical ITR regions were shown in the Orf virus and Bovine Papular Stomatitis virus genomes (47).

### Identifying Inverted Terminal Repeats (ITRs)

We also identified Inverted Terminal Repeats (ITRs) by filtering out all inverted repeats that do not occur at the start or end of genomic sequences. Using a Python script, we identified ITRs in 295 viral genomes. We listed the viruses with the longest inverted terminal repeats below:

## Conclusions

It can be inferred from the pilot study that BLMT consistently obtained the highest total number of inverted repeats in all of the sample viruses in comparison to EMBOSS, IUPACpal, detectIR, and DNA Analyzer. On the other hand, EMBOSS, DNA Analyzer, and IUPACpal almost equally identified the same number of inverted repeats. The latter is only a few sequences above EMBOSS in terms of the number of identified inverted repeats. DetectIR, however, found no IRs common to other tools in all of the viruses, which suggests that it completely failed to detect other existing IRs within the genomes. Based on the data, BLMT showed the greatest number of unique sequences. This proves that BLMT is a rather novel bioinformatics tool that can challenge and surpass existing inverted repeat detection tools in terms of accurately identifying imperfect inverted repeats.

Our analysis also provides a rigorous analysis of inverted repeats (IRs) in viral genomes, shedding light on their structural characteristics and implications for viral biology. By examining statistical parameters such as the mean and standard deviation of IR lengths, the normalized frequency of IRs, the positions of IRs within genomes, and their correlation with viral hosts, we have achieved a comprehensive understanding of IRs in viral genomes. The analysis of mean IR lengths revealed an average range of 10 to 11 across the examined viruses. Notably, variations in IR lengths were observed among different virus categories, with viruses infecting human, invertebrate, and vertebrate hosts displaying higher standard deviations, indicating greater heterogeneity in IR lengths within these host groups. Conversely, viruses infecting bacteria, fungi, and land plants exhibited lower standard deviations, suggesting a more consistent pattern of IR lengths. The examination of the normalized frequency of IRs provides insights on their prevalence in viral genomes. Bacteria-hosted viruses exhibited a higher frequency of IRs, while viruses infecting fungi and land plants displayed relatively lower frequencies. Remarkably, viruses infecting human, invertebrate, and vertebrate hosts demonstrated a similar mean and standard deviation of normalized IR frequencies, suggesting potential evolutionary similarities and functional implications among these host species. The analysis of IR positions within viral genomes revealed a preference for their occurrence in the latter half of the genome, with a concentration towards the middle region rather than the start or end. This positional distribution highlights the potential functional significance of IRs in viral gene expression and replication processes. Furthermore, our identification of specific viruses with notable characteristics, such as high average IR lengths and extended inverted terminal repeats (ITRs), provides further insights into the genomic structures of these viral species. Species of herpesviruses, densoviruses, and parvoviruses exhibited significant variations in IR lengths, highlighting the diversity within these viral families. Bacteriophages and poxviruses also demonstrated considerable lengths of ITRs, suggesting their importance in viral replication and genetic stability. Our findings contribute to the understanding of viral genome organization and the prevalence of IRs across different virus categories. The insights gained have implications for viral replication, evolution, and host-virus interactions. The comprehensive analysis of IRs in viral genomes enhances our knowledge of viral biology and provides potential avenues for the development of targeted antiviral strategies. In summary, this study highlights the significance of IRs as crucial genomic features in viruses and underscores their diverse characteristics within different virus categories. The examination of statistical parameters and their correlations with viral hosts has advanced our understanding of IRs and their potential functional roles in viral genomes. Further investigations into the functional significance of IRs and their implications for viral pathogenesis will deepen our knowledge of viral biology and inform the development of novel therapeutic approaches.

## Author Contributions

MKG and MSh conceptualized the problem to analyze palindromes in viral genomes. MKG had previously developed the Biological Language Modeling Toolkit, and here developed additional features for viral palindrome identification. MSe, JG and GMR were undergraduate students in three different geographic locations (India, Qatar, Philippines) when this work carried out; MSe and JG received exclusively remote guidance (video calls) from MKG. JG carried out literature reviews. MSe carried out a comparison of different tools with inputs from GMR and MKG. JG carried out post-processing of results on viral families. Manuscript has read and approved by all authors.

